# Identification of perrhenate-binding peptides by phage display

**DOI:** 10.1101/2025.08.12.669540

**Authors:** Samuel Takyi, Mark Aldren M. Feliciano, Sooriyage Salika Dulanjali, Brian Gold, Mark C. Walker

## Abstract

Pertechnetate is the most stable and highly environmentally mobile form of the radioactive element technetium. To explore the ability of peptides to remove this molecule from aqueous solutions, a phage display library was biopanned against immobilized perrhenate, a nonradioactive analog of pertechnetate. Six unique peptides were identified from the screen and their ability to bind perrhenate free in solution and immobilized on a solid support were explored. It was found that the peptides, particularly when immobilized, were able to remove perrhenate from aqueous solutions, and despite not being screened for selectivity, demonstrated some preference for perrhenate over other anions such as chromate and nitrite. These results demonstrate the feasibility of using engineered biological systems for remediation of pertechnetate in the environment.

## Introduction

99-Technetium is a radioactive nuclide present in nuclear waste streams with a relatively long half-life of 2.13 × 10^5^ years. The most stable chemical form is the pertechnetate anion ([TcO_4_]^-^), which is highly soluble in water and environmentally mobile, raising environmental and ecological concerns should it be released from those streams. Additionally, pertechnetate is volatile enough that only a fraction is incorporated during vitrification to immobilize nuclear waste into borosilicate glass to immobilize it [1]. These properties make technetium a priority for remediation and containment. Current approaches to immobilize pertechnetate involve materials such as organic polymers [2-4], covalent organic frameworks [5], layer double hydroxides [6], among others, or reduction of pertechnetate to an much less soluble form of technetium [7, 8]. These separations are challenging due in part to the presence of other anions in nuclear waste streams at much higher concentrations than pertechnetate [9, 10]. A potential approach to address this challenge is to utilize biological systems that can have exquisite selectivity for their target compounds [11].

Phage display is a technique that was originally developed for affinity maturation of antibodies [12]. While multiple platforms have been developed to anchor proteins or peptides to the surface of phage particles, a common approach is to genetically fuse genes encoding the library members to the gene encoding protein III of the M13 phage. This technique allows up to five copies of the same protein or peptide to be anchored to the surface of a phage particle. The phage particle also contains the DNA that encoded the library member, attaching the protein or peptide to the genetic information that encoded it. Phage are then mixed with the target, target binders are then pulled down, and the protein or peptide that mediates the interaction between the target and the phage can be identified by sequencing the genetic material that encoded it. This process allows for the screening of up to 10^9^ different amino acids sequences for the ability to bind to a target. Phage display has been used to identify peptides with the ability to bind inorganic matter such as GaAs [13], ZnS [14], SiO_2_ [15], TiO_2_ [16], and many other examples [17].

To explore the potential of using peptides to selectively remove pertechnetate from nuclear waste streams we sought to identify peptides that were able to bind to perrhenate, a nonradioactive surrogate of pertechnetate often used in laboratory studies.

## Materials and Methods

### Commercial materials

Luria-Bertani (LB) Broth Miller, LB Agar Miller, and glycerol were purchased from EMD Biosciences (Darmstadt, Germany). Isopropyl-b-D-thiogalactopyranoside (IPTG), 5-bromo-4-chloro-3-indolyl-beta-D-galacto-pyranoside (X-gal), tris(hydroxymethyl)aminomethane hydrochloride (Tris-HCl), sodium chloride, 4-(2-hydroxyethyl)-1-piperazineethanesulfonic acid (HEPES), and tetracycline, were purchased from Fisher Scientific (Pittsburgh, United States). Dowex 1×8 200-400 (Cl), Phosphate Buffered Saline, and sodium perrhenate were purchased from Sigma-Aldrich (St. Louis, United States). All 20 amino acids were purchased from Combi-Blocks, Inc. (San Diego, CA). All solvents were purchased from VWR International, LLC (Radnor, United States), and the organic solvents used were of analytical grade. The Phage Displayed Peptide library (Ph.D. 7) was procured from New England Biolabs (Ipswich, United States). UV-Vis data were recorded on Cary 60 UV-Vis spectrometer (Agilent Technologies, Santa Clara, United States). Purification was done using DIONEX UltiMate 3000 HPLC (Thermo Fisher Scientific, Waltham, United States).

### Perrhenate immobilization

Experiments were performed as previously described [18]. 20 mg Dowex 1×8 200-400 (Cl) was packed in a glass pipette stoppered with cotton wool. 10 ml of normal saline was passed through the resin column followed by 5 ml of water. To determine the loading capacity of the packed column, sodium perrhenate stock solution of 10 mg/ml was prepared. 2 mg of NaReO_4_ was loaded onto the resin. Three buffers (HEPES, HEPES saline, and Phosphate Buffer Saline; 20 mM, pH 7.4) were used to determine the ideal buffer for the experiment. Each of these buffers were passed over a NaReO_4_ loaded resin column and the amount of perrhenate displaced was determined using a Cary 60 UV spectrophotometer. Absorbances at 225 nm were compared to a standard curve to determine the quantity of perrhenate present in the eluent.

### Panning of phage library

Biopanning experiments were performed according to manufacturer’s protocol. Briefly, approximately 5.76 × 10^12^ phage were passed over a NaReO_4_ loaded resin column. The column was washed with 5 mL of wash buffer (20 mM of HEPES buffer at pH 7.4). 1 mL of 10 mg/ml NaReO4 in wash buffer was used to elute the retained phage. The eluted phage were amplified by adding them to a 20 mL culture in a 250 mL Erlenmeyer flask inoculated with 200 µL of an overnight culture of *E. coli* K12 ER2738 (New England Biolabs) culture. This culture grown at 37°C with shaking at 250 rpm for 4 - 5 hrs. Cells were removed by centrifugation at 4,500 × g for 10 min and the supernatant was transferred to a fresh tube. 16 ml of supernatant was transferred to a new tube and 4 mL of 2.5 M NaCl/20 % PEG-8000 (w/v) was added and briefly mixed. The phage were precipitated overnight at 4°C. The phage were collected by centrifugation at 12,000 × g for 15 min at 4°C. The supernatant was discarded, and the pellet was resuspended in 1 mL of Tris Buffered Saline at pH 8.0. The resuspended phage were transferred to an Eppendorf tube and spun briefly to remove any cell debris. The supernatant was transferred to a fresh tube and the phage were precipitated a second time by adding 200 µL of 2.5 M NaCl/20% PEG-8000 and incubating on ice for 15-60 min. The phage were collected by centrifugation at 16,000 × g in a benchtop centrifuge at room temperature for 10 min. The supernatant was discarded, the tube was spun down briefly, and the remaining supernatant was removed with pipette and the pellet was resuspended in 200 µL Tris Buffer Saline pH of 8.0. These amplified phages were subjected to another round of biopanning and the process was repeated for a total of six rounds of biopanning.

### Tittering of eluted phage

Tittering of the phage in the eluate was performed according to the manufacturer’s protocol [19]. Briefly, a colony of *E. coli* ER2738* cells grown overnight at 37°C on an LB plate supplemented with 20 mg/mL of tetracycline was used to inoculate 5 mL of LB that was subsequently incubated with shaking for 4 – 8 hours until the culture reached mid-log phase (OD_600_ ∼ 0.5). At the same time, serial dilutions from 1:10 to 1:10,000 of the phage containing eluate were prepared. 10 μL of each serial dilution was added to 200 μL of the *E. coli* culture, mixed, and incubated at room temperature for 1 – 5 min. These mixtures were added to molten Top Agar, mixed, and poured onto LB plates containing 0.05 g/mL IPTG and 0.04 g/mL Xgal. The plates were gently tilted and rotated to spread the Top Agar, allowed to cool at room temperature for 5 min, and then incubated overnight at 37°C. Plates that had approximately 100 plaques were counted and the plaque-forming units per volume were determined.

### Phagemid sequencing

After the final round of biopanning, 20 plaques were randomly picked and amplified. The resulting phage pellet was suspended thoroughly in 100 μL of Iodide Buffer (10 mM Tris-HCl, 1 mM EDTA, 4 M sodium iodide, pH 8.0) by vigorously tapping the tube. Single-stranded phage DNA was precipitated by the addition of ethanol (250 μL) followed by incubation for 10 – 20 minutes at room temperature. Precipitated DNA was collected by centrifugation at 20,000 × g for 10 minutes. The pellet was washed with 0.5 mL of cold (−20°C) 70% ethanol, re-spun, discarded the supernatant, and the pellet was briefly dried under vacuum. The dried pellet was dissolved in 30 μL of TE buffer (10 mM Tris-HCl, 1 mM EDTA, pH 8.0) and analyzed by Sanger sequencing (Azenta Life Sciences, South Plainfield, United States) using the -96gIII sequencing primer (5’-CCCTCATAGTTAGCGTAACG-3’).

### Synthesis of free peptides

Peptides were synthesized using the solid-phase peptide synthesis (SPPS) method and the Fmoc strategy [20]. As the identified peptides were anchored to the phage at the c-terminal end (i.e. did not have a free c-terminus), the peptides were synthesized with a c-terminal amide. 150 mg of Rink Amide resin was used for a scale of 0.05 mmol scale synthesis. The resin was transferred to a peptide reactor (Biotage LLC, Uppsala, Sweden) and resin was swollen in 3 – 5 mL DCM for 30 minutes with agitation. The resin was washed three times with DMF prior to the Fmoc cleavage from the resin using 40% piperidine in DMF for 10 minutes. Based on the amount of resin, 4:3.6:8 equivalences of the amino acids, HBTU and DIPEA respectively were used to achieve 100% loading of the resin. Amino acids were activated by the addition of HBTU and DIPEA in a glass scintillation vial in 2 mL of DMF. After cleavage of the Fmoc from the resin was complete, a vacuum pump was used to drain the 40% piperidine. The resin was washed three times with DMF and the amino acid/HBTU/DIPEA solution was added to the resin and rocked at room temperature for 1 hour. After 1 hour, a vacuum pump was used to drain the amino acid solution. The Fmoc on the N-termini was cleaved using 40% piperidine in DMF and the loaded resin was washed with DCM 3x and DMF 3x. This cycle was repeated for the remaining amino acids. After the last amino acid was coupled and Fmoc deprotected, the loaded resin was dried under high vacuum overnight. The dried resin was transferred into a falcon tube and 95:2.5:2.5 TFA: H_2_O: TIPS were added to the resin and left to rock for 1.5-2 hours. The resulting solution was concentrated by blowing air over TFA solution, and the peptides were precipitated with cold anhydrous ethyl ether. The peptides were collected by centrifugation (8,000 × g for 3 minutes), the supernatant was decanted, and the pelleted precipitate was left to dry. The dried precipitate was dissolved in 50:50 ACN/H_2_O and the resin were removed by filtration. The filtrate was frozen using liquid nitrogen and lyophilized using the FreeZone Lyophilizer (Labconco Corporation, Kansas City, United States). The dried crude peptide was purified using the 1260 Infinity II HPLC (Agilent Technologies). The peptides were purified using a Macherey-Nagel (Duren, Germany) C18 column with dimensions of 250 mm × 10.0 mm and 10 µm particle size. A gradient from 98% water with 0.1% TFA and 2% acetonitrile with 0.1% TFA to 100% acetonitrile with 0.1% TFA over 60 min was used to elute the peptides. Fractions containing the desired peptides were identified by mass spectrometry with a Waters QDa single quadruple mass spectrometer (Waters Corporation, Milford, United States).

### Synthesis of immobilized peptides

Immobilized peptides were synthesized using Tentagel S NH_2_ (VWR) as solid support, using Fmoc SPPS protocol. 200 mg of Tentagel S NH_2_ resin was used to achieve a loading capacity of 0.04 mmol scale for the synthesis. The resin was added to a peptide reactor. The resin was swollen in DCM for 30 mins and washed three times with 5 mL of DMF. The Fmoc was cleaved from the resin three times with 5 mL of 40% piperidine in DMF for 10 minutes. The activation of the amino acids was carried out as described for the free peptides. After the first amino acid was coupled to the resin, the remaining free amines on the resin were capped using 10% acetic anhydride in DMF and DIPEA. The Fmoc on the N-termini of the first coupled amino acid was deprotected using 40% Piperidine in DMF and washed with DCM 3x and DMF 3x. The cycle was repeated for the remaining amino acids and the acid labile protecting groups were deprotected using 95:2.5:2.5 TFA: H_2_O: TIPS respectively for 1.5 - 2 hours. The loaded resin was dried overnight using a high vacuum and the mass of the peptide was determined.

### Procedure for NMR Titrations

Experiments were performed as previously described [21]. All of the reagents were weighted separately on an analytical balance (readability to 0.0001 mg) in 1.5 mL Eppendorf tubes. D_2_O-d_2_ was used as the solvent for NMR titration. A solution of 2 mM peptide was prepared for the NMR analysis. 200 mM solution of NaReO_4_ in 2 mM peptide solution was prepared to avoid the diluting the peptide during the titration. The titration was performed by adding aliquots (20 µL) of the perrhenate solution to the peptide solution (400 µL) and recording the ^1^H NMR spectrum following each addition of perrhenate solution. The temperature inside the NMR probe was held constant at 25^?^C. Association constants were calculated based on the changes in the chemical shifts of the most affected protons of the peptide.

### Retention of metal anions by immobilized peptides

The mass of corresponding to 4.3 mg of each resin-bound peptide was loaded in a 1 mL disposable syringe plugged with cotton to retain beads. The resin was washed five times with 1 mL of wash buffer (HEPES buffer, 20 mM, pH 7.4) to swell and equilibrate the resin as well as wash off any remaining impurities. Solutions of 2 mg/mL NaReO_4_, NaNO_2_, Na_2_CrO_4_, and NaI were prepared in the wash buffer. 500 µL of the metal anion solutions were incubated with the resin-bound peptide column for 15 minutes with shaking at room temperature. The remaining solution was drained into scintillation vial for UV-Vis analysis. The resin column was washed five times with 1 mL of Milli-Q water. These washes were collected to assess metal loss during washing.

### Competition with sodium chloride

Perrhenate was bound to the resin-bound peptide columns as described above. 500 µL of NaCl (2 mg/mL in HEPES, 20 mM, pH 7.4) was incubated with the resin containing the retained NaReO_4_ for 15 minutes with shaking. The solution was drained after 15 minutes to assess the displaced NaReO_4_.

### Quantification by UV-Vis spectrometry

Metal anion concentrations in the flowthrough and wash fractions were quantified using UV–visible spectroscopy. Absorbance measurements were recorded using a Cary 60 UV-Vis spectrometer at the appropriate wavelength for each metal anion (372 nm for chromate, 230 nm for perrhenate, 235 nm for iodide and nitrite). Standard curves were generated from serial dilutions of metal salt solutions prepared in the wash buffer, with linearity confirmed over the relevant concentration range. The concentration of metal anions in each fraction was determined by interpolating from the standard curve. The % retention of metal ions by the peptide-functionalized resin was calculated using the following equation:

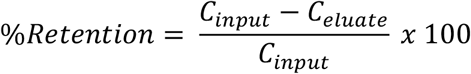

Where *C*_*input*_ is the concentration of metal in the input solution, and *C*_*eluate*_ is the combined concentration of metal in the flowthrough and wash fractions. All measurements were performed in triplicate.

## Results and Discussion

The random Ph.D.-7 phage library was screened to identify peptide sequences that bind specifically to perrhenate. The intended application of these peptides was ultimately to be utilized in the sequestration of pertechnetate from off-gas condensate of low-activity nuclear waste. The unmodified phage screening identified major hit peptides in this study.

### Screening of Phage Library

An increase in the number of phage eluted was observed after the third round of biopanning, but sequencing select phages did not reveal an enrichment in any particular peptide sequences. Therefore, three more rounds of biopanning were carried out (*Figure 1*). Phagemids from 20 randomly selected plaques were sequenced, with 12 of those clones providing unambiguous sequence data. These 12 clones represented 9 unique peptide sequences (*Figure 1*). None of the identified peptides contained a negatively charged residue, suggesting the phage were not being enriched for the ability to bind to the anion exchange column. Furthermore, all of the peptides contained at least one positively charged residue, which could be important for binding to the perrhenate anion. Out of the 9 unique peptides, 5 began with the same sequence (KMI). These five peptides were selected for further characterization (ST1 – ST5) as well as another peptide that appeared twice among the sequenced clones (ST6).

**Figure 1.**
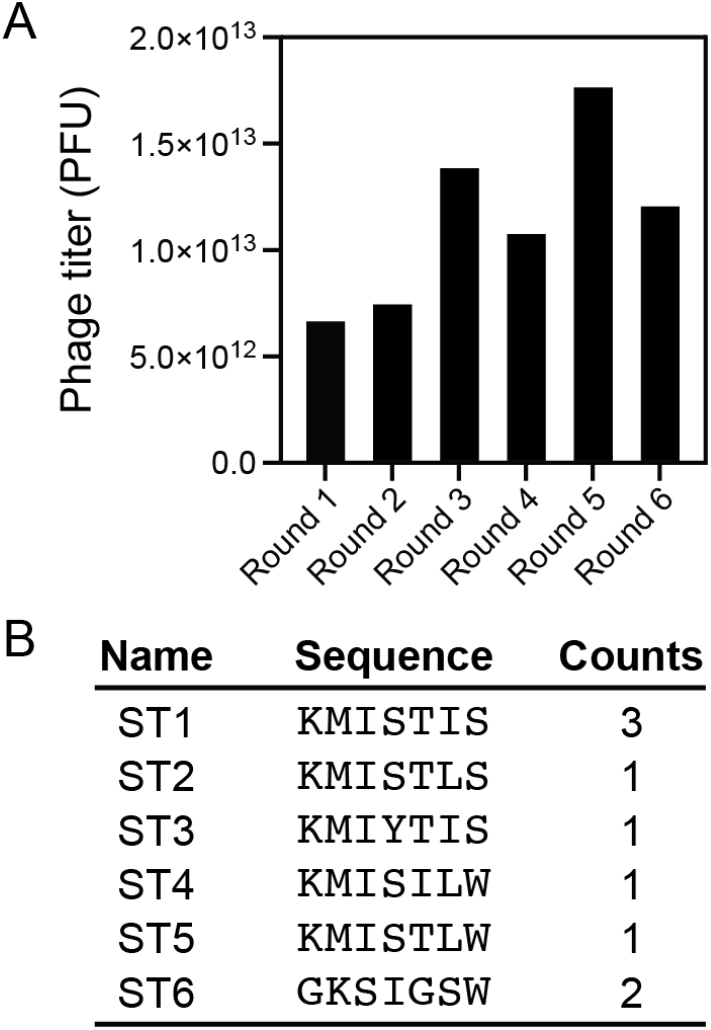
Biopanning against immobilized perrhenate. A) Titer of phages eluted from column with immobilized perrhenate. B) Peptide sequences identified after 6 rounds of selection. Counts are the number of times the gene encoding the given peptide was sequenced.

### Perrhenate binding

To confirm that the peptides identified by biopanning bound perrhenate, these peptides were synthesized (*Supplementary Figures S1 – S6*) and examined by ^1^H NMR with increasing concentrations of perrhenate (*Supplementary Figures S7 – S12*). The spectra of ST1, ST2, ST3, and ST5 in D2O revealed a downfield shift of the protons of the thioether methyl group of the methionine side chain as the concentration of perrhenate increased. The spectra of ST4 revealed the appearance of a new peak and disappearance of the methionine peaks as the concentration of perrhenate increased. These alterations in the spectra of the peptides may indicate the formation of a complex, however, the peptides could not be saturated by increasing concentrations of perrhenate. To estimate the affinity of the peptides for perrhenate, the changes in chemical shift were fit to the Benesi-Hildebrand equation (Table 1). While none of the peptides exhibited particularly strong affinity for perrhenate, in phage display multiple copies of a peptide are located close together and the avidity of the phage particle for the perrhenate could be higher than the affinity.

**Table 1.** Affinity of peptides for perrhenate determined by NMR.

| Peptide | Association constant (M<sup>-1</sup>) |
| --- | --- |
| ST1 | 17 ± 1 |
| ST2 | 11.8 ± 0.4 |
| ST3 | 21 ± 1 |
| ST4 | 27 ± 2 |
| ST5 | 10.5 ± 0.2 |
| ST6 | 3.3 ± 0.1 |

Interestingly, as ST4 was incubated with perrhenate a precipitate formed. As precipitation of perrhenate could be a potential approach for its remediation, further characterization was performed. A scrambled peptide with the sequence LMIWKSI was synthesized and incubated with perrhenate. This scrambled peptide did not result in the formation of a precipitate, suggesting that the sequence of ST4, not simply the presence of the same functional groups, is responsible for the formation of the precipitate. Additionally, truncated peptides with the sequences KMIS (*Supplementary Figure S13*) and KMIY (*Supplementary Figure S13*) were synthesized and incubated with perrhenate. Again, no precipitation formed, suggesting the c-terminal residues of ST4 play a role in the formation of the precipitate.

### Binding and selectivity of immobilized peptides

To test the ability of these peptides to remove perrhenate from solution we synthesized the peptides immobilized on a solid support. Approximately 5 *μ*moles of the peptides were incubated with approximately 3.7 *μ*moles (1 mg) of perrhenate in a buffer solution for 15 minutes. All of the immobilized peptides were able to remove the majority of the perrhenate from the buffer solution (*Figure 2*). ST5 bound the largest fraction of perrhenate, at 93%, while ST1 retained the least, with 86%. Washing the immobilized peptides with water did not result in the elution of a detectable amount of perrhenate. To explore the ability of the peptides to maintain an interaction with perrhenate in the presence of other ions, they were washed with 34 mM sodium chloride. This wash resulted in the removal of some, but not all, perrhenate from all of the peptides (*Figure 2*). In addition to binding the largest amount of perrhenate, ST5 lost the largest amount in the presence of sodium chloride, at about 33% of what it had bound. ST4 lost the smallest amount of perrhenate in the presence of sodium chloride, with about 17% of what it had bound. This ability to retain perrhenate may be due to the same process that led to ST4 precipitating in its presence.

**Figure 2.**
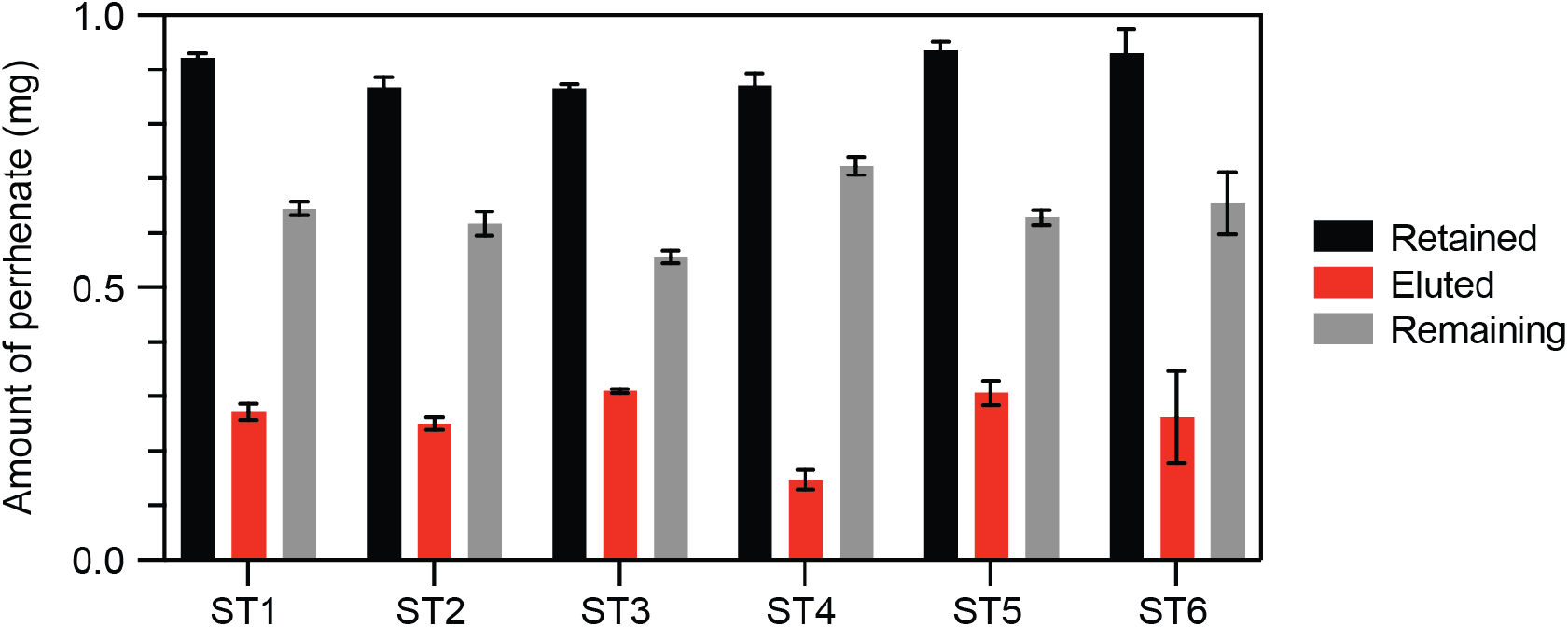
Retention of perrhenate in the presence of sodium chloride. The amount of perrhenate retained by peptides immobilized on resin in the absence of NaCl (black), the amount of perrhenate eluted by washing with NaCl (red), and the amount of perrhenate remaining on the column following the NaCl wash (grey). The error bars represent the standard deviation of three replicates.

While the identified peptides were selected for their ability to bind perrhenate, their ability to bind other anions was explored. All of the peptides were able to bind chromate, nitrite, and iodide to some extent (*Figure 3)*. However, the amount of chromate and nitrite the peptides were able to bind was approximately 20 – 25% of the amount of perrhenate they were able to bind. Additionally, the amount of iodide the peptides were able to bind was around 50% of the amount of perrhenate. These results suggest that the peptides have some level of specificity for perrhenate despite not having been screened for the inability to bind other anions.

**Figure 3.**
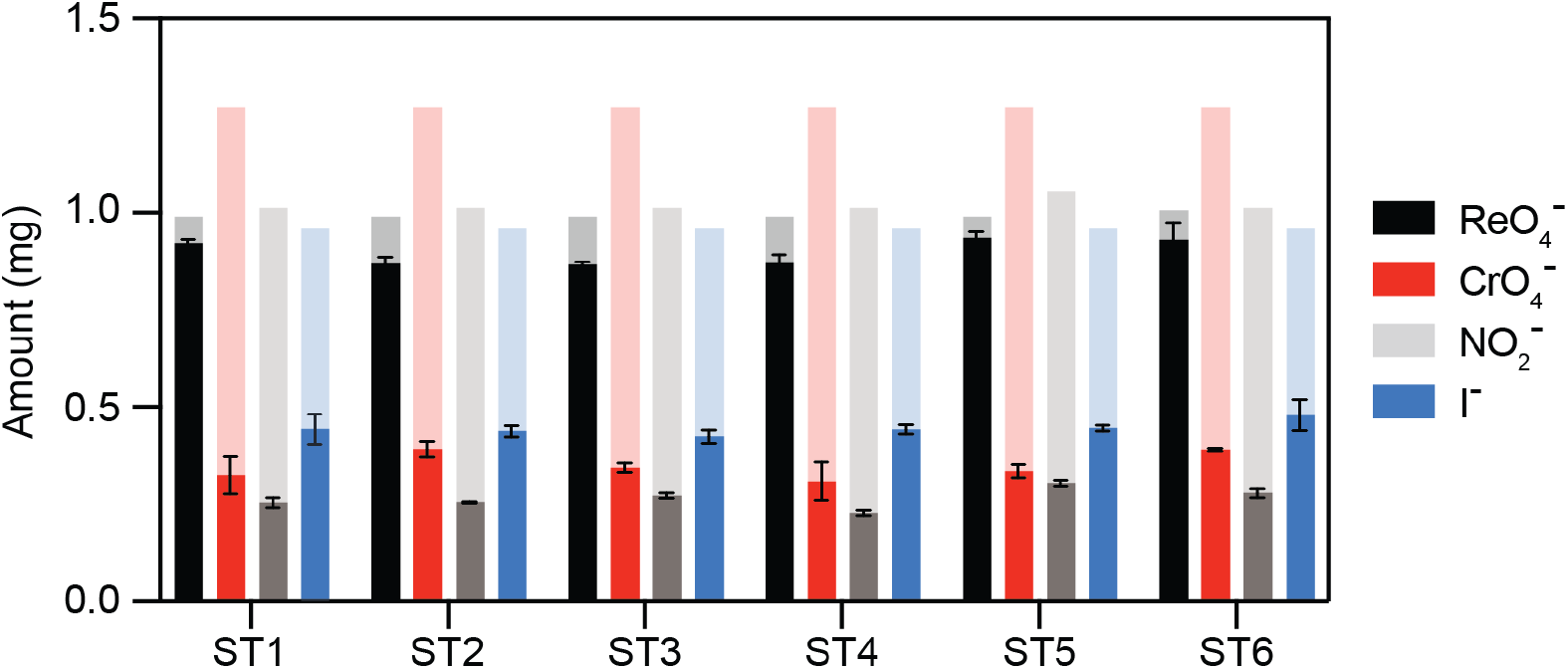
Retention of other anions by immobilized peptides. The amount of select anions loaded (light colors) onto immobilized peptides is compared to the amount retained by those peptides (solid colors). The error bars represent the standard deviation of three replicates.

## Conclusion

In this study we sought to explore the potential of using biological materials to remove perrhenate from aqueous solutions. We used phage display to identify several peptides that were able to interact with perrhenate immobilized on anion exchange resin. We then found that six of these peptides, in their soluble form, were able to interact with free perrhenate with modest association constants. Furthermore, we were able to show that when the peptides were immobilized on resin, that resin was able to remove the majority of perrhenate in buffered solution and was somewhat selective for perrhenate over other anions such as chromate. These studies suggest, while further optimization is required, that biological materials can be engineered to immobilize perrhenate. Future alterations of the peptides such as cyclization to lock in a certain conformation or inclusion of nonproteinogenic amino acids could result in generation of just such materials.

## Notes

### Competing Interest Statement

The authors have declared no competing interest.

### Funding

- United States Department of Energy

