## Supplementary figures S1-S15 for "Identification of perrhenate-binding peptides by phage display"

**Supplementary Figure S1.** Characterization of ST1. A) High resolution mass spectrum. [M+H] expected m/z: 778.4491, Δppm: 3. B) HPLC chromatogram of purified ST1.

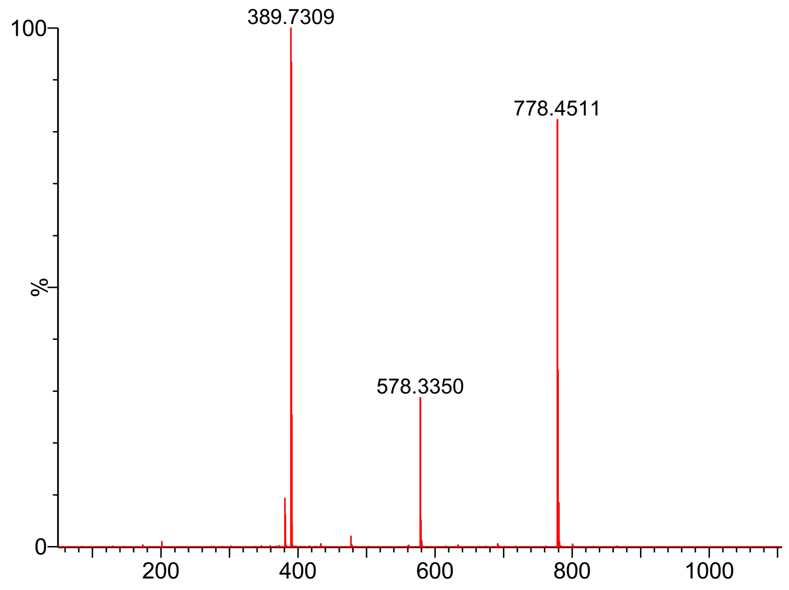


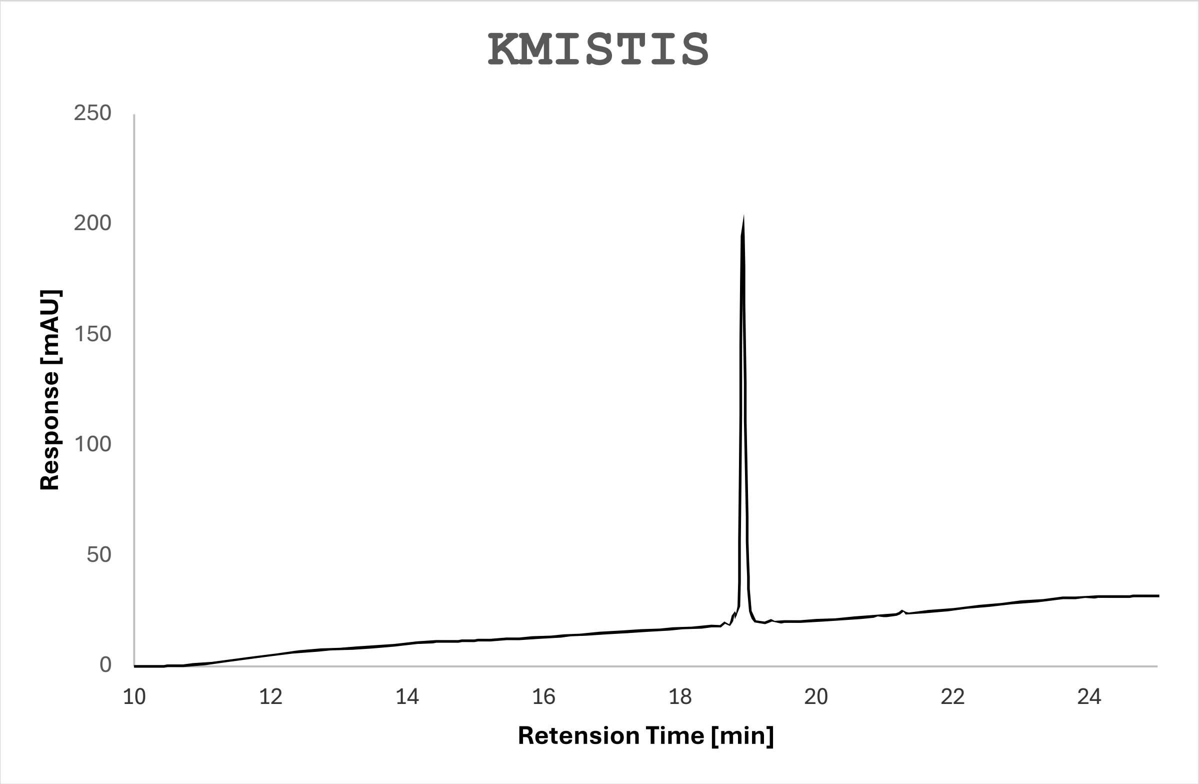

**Supplementary Figure S2.** Characterization of ST2. A) High resolution mass spectrum. [M+H] expected m/z: 778.4491, Δppm: 3. B) HPLC chromatogram of purified ST2.

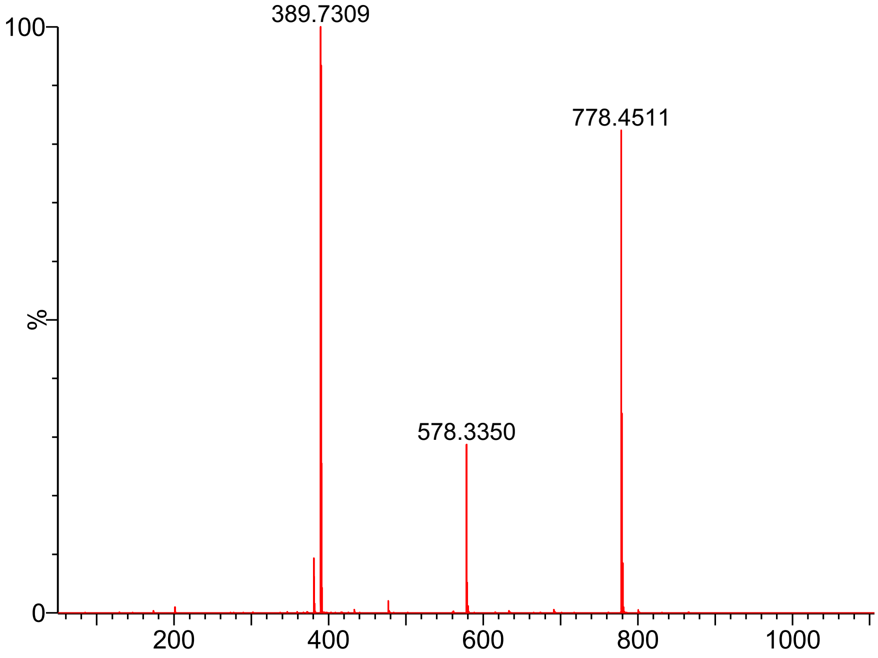


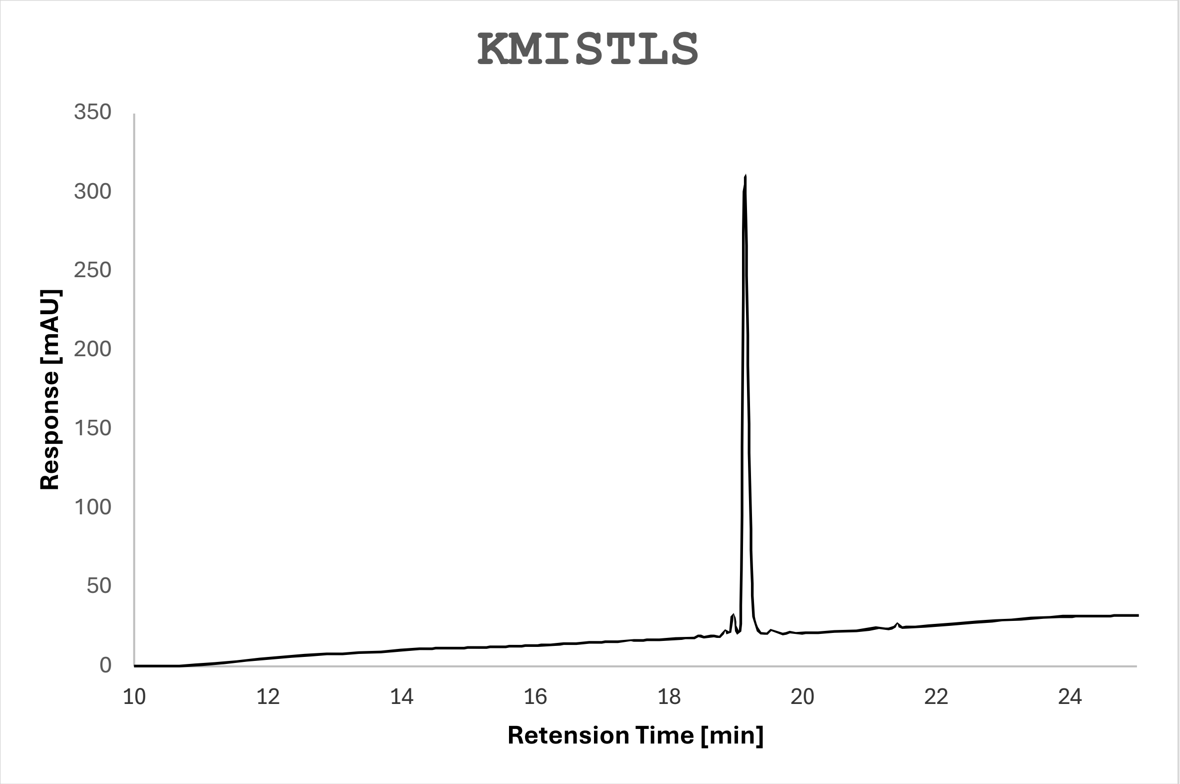

**Supplementary Figure S3.** Characterization of ST3. A) High resolution mass spectrum. [M+H] expected m/z:  854.4804, Δppm: 11. B) HPLC chromatogram of purified ST3.

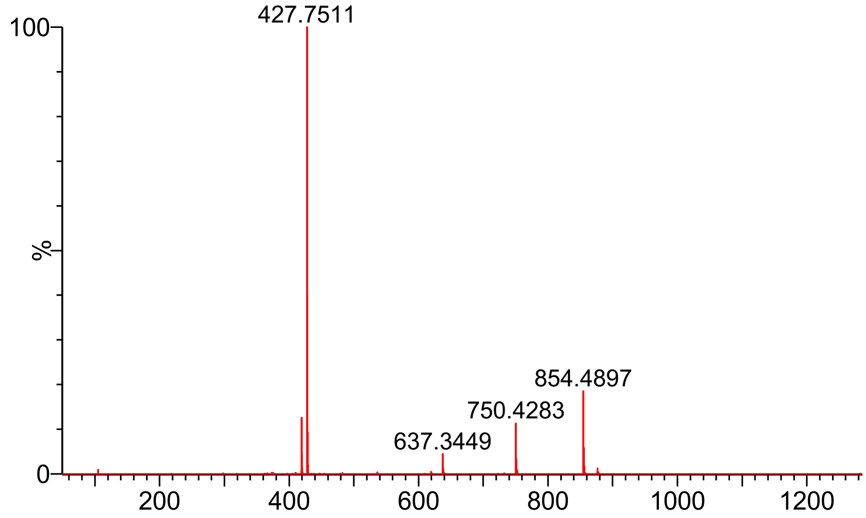


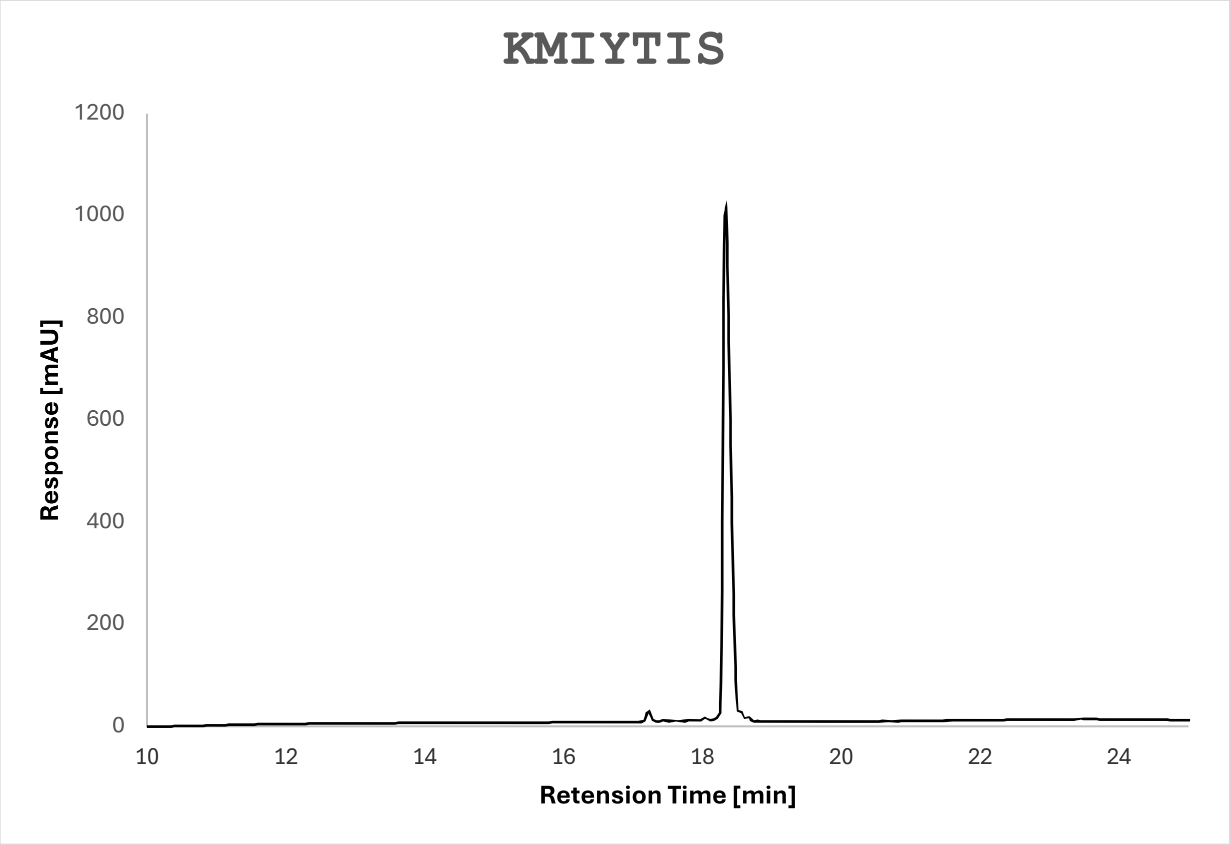

**Supplementary Figure S4.** Characterization of ST4. A) High resolution mass spectrum. [M+H] expected m/z:  889.5328, Δppm: 12. B) HPLC chromatogram of purified ST4.

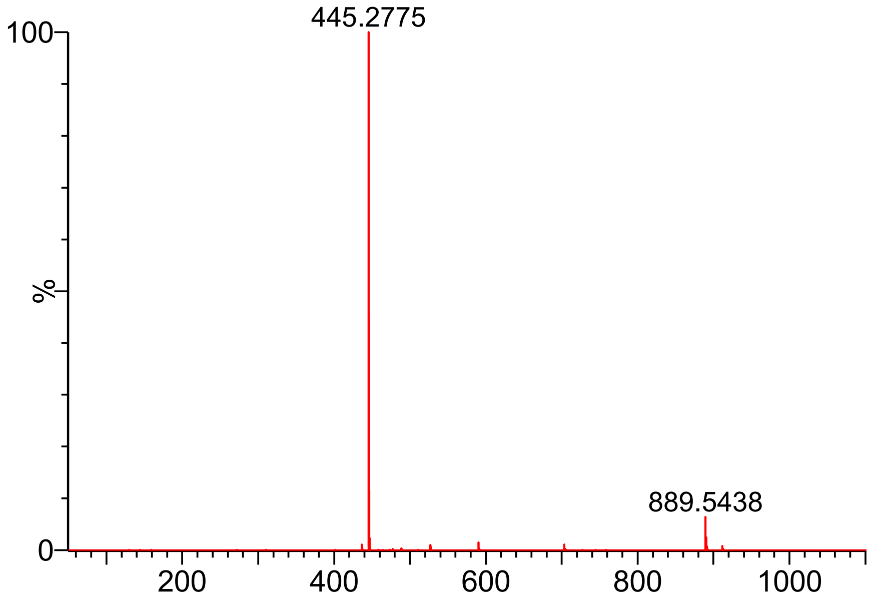


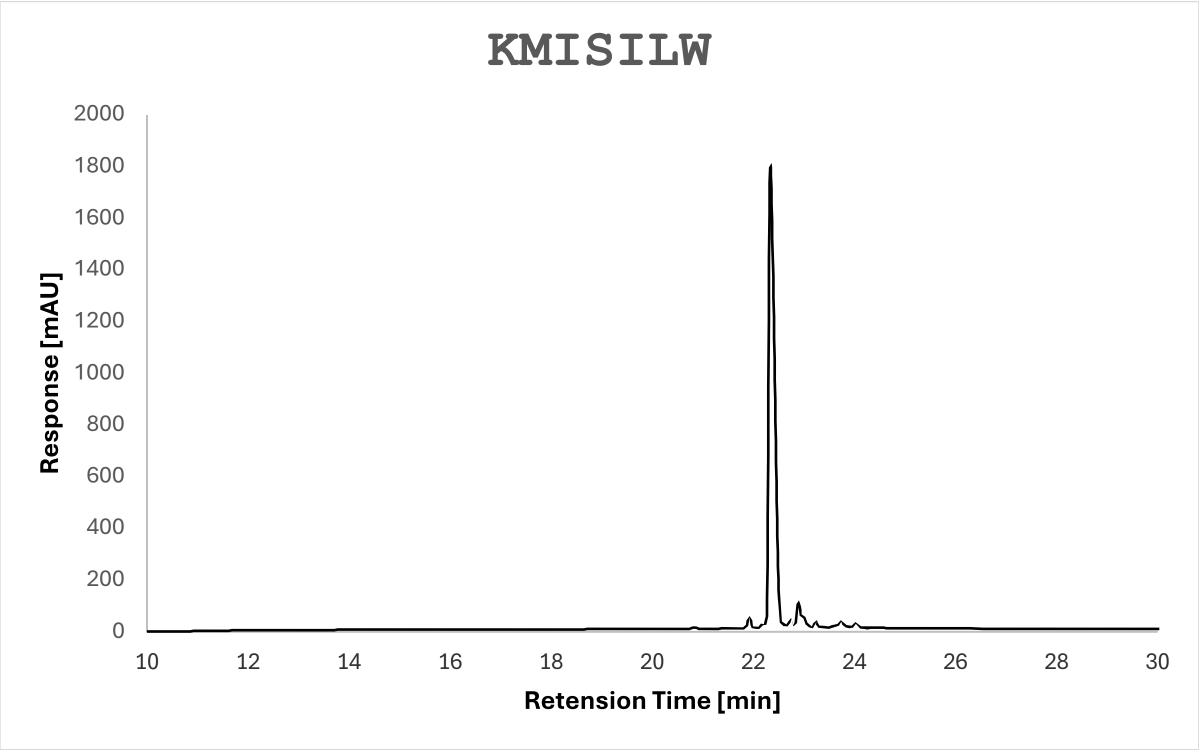

**Supplementary Figure S5.** Characterization of ST5. A) High resolution mass spectrum. [M+H] expected m/z: 877.4964, Δppm: 13. B) HPLC chromatogram of purified ST5.

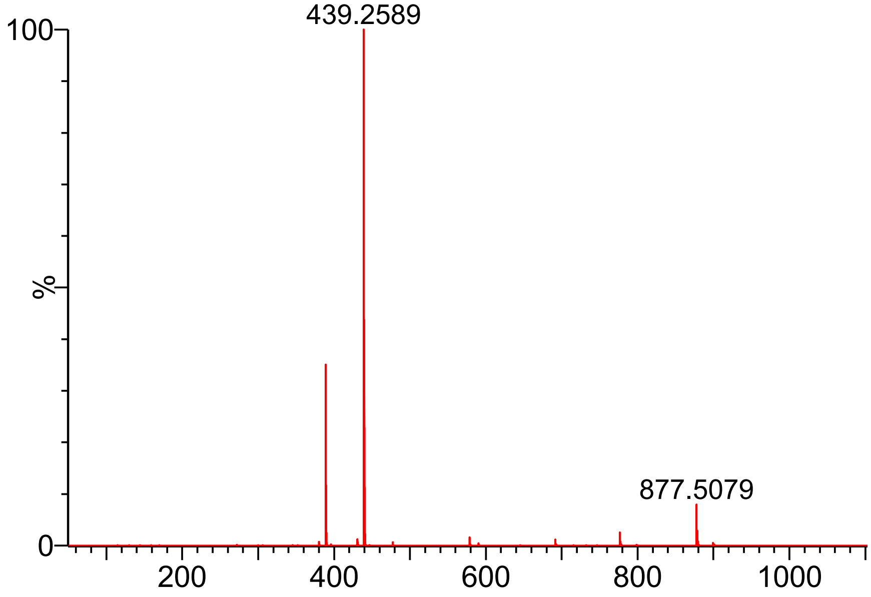


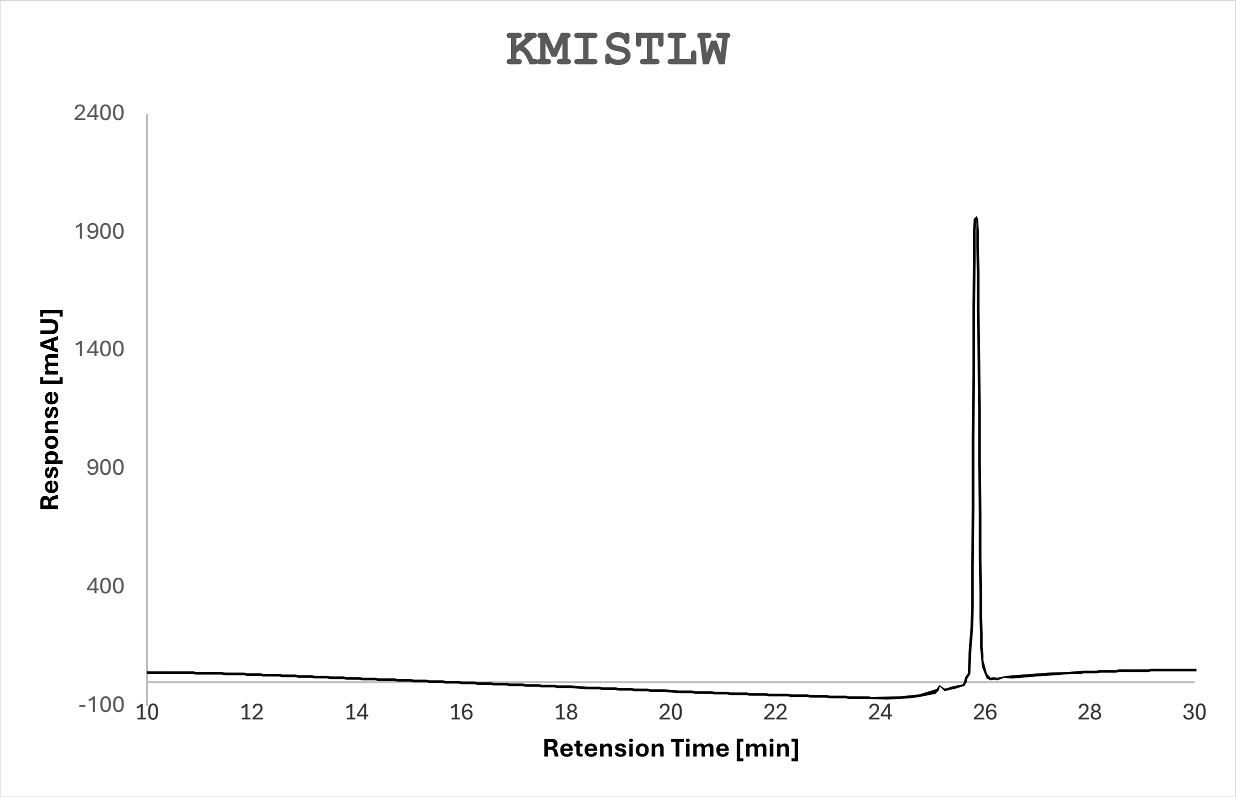

**Supplementary Figure S6.** Characterization of ST6. A) High resolution mass spectrum. [M+H] expected m/z: 733.3991, Δppm: 16. B) HPLC chromatogram of purified ST6.

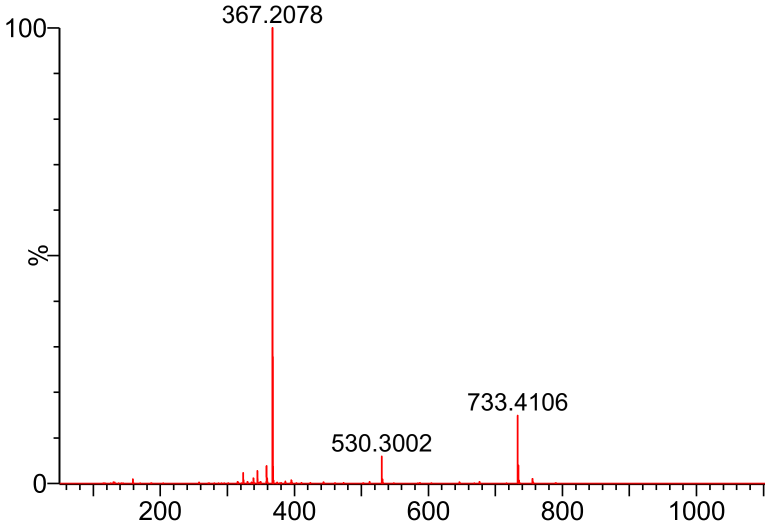


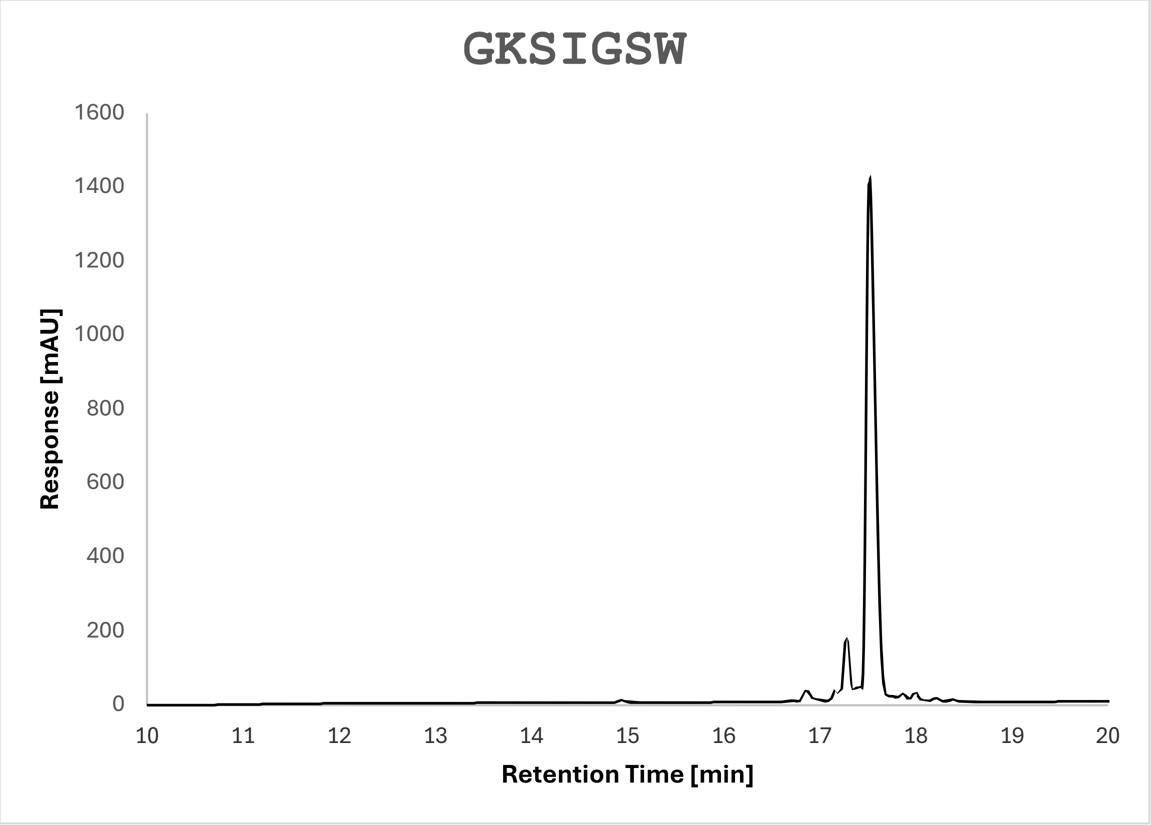

**Supplementary Figure S7.** NMR titration of ST1.

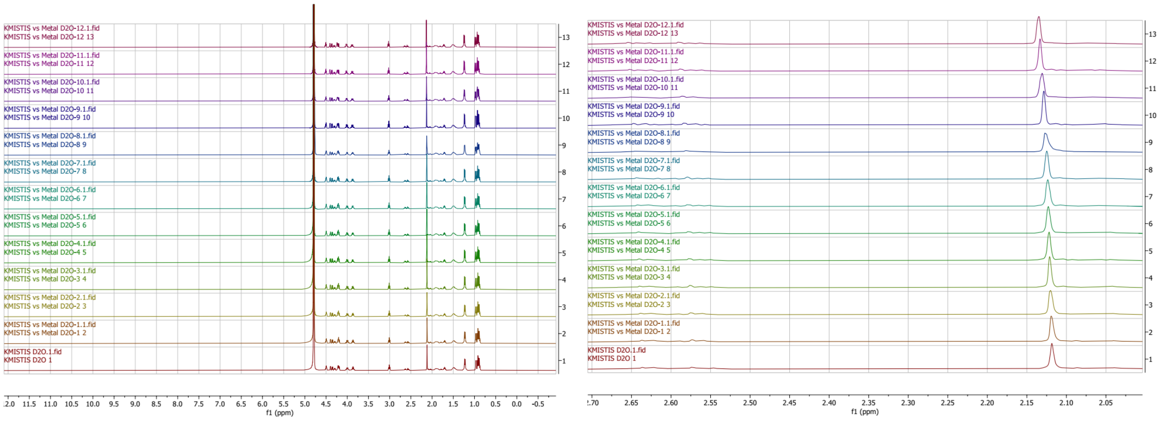


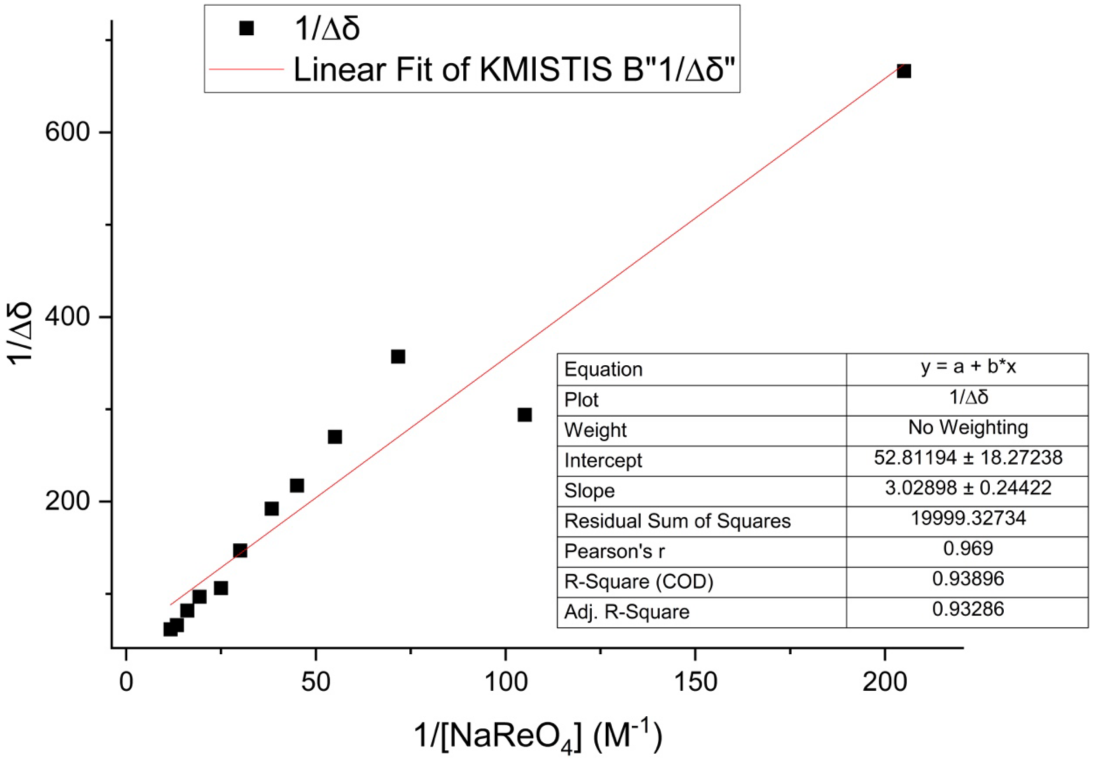

**Supplementary Figure S8.** NMR titration of ST2.

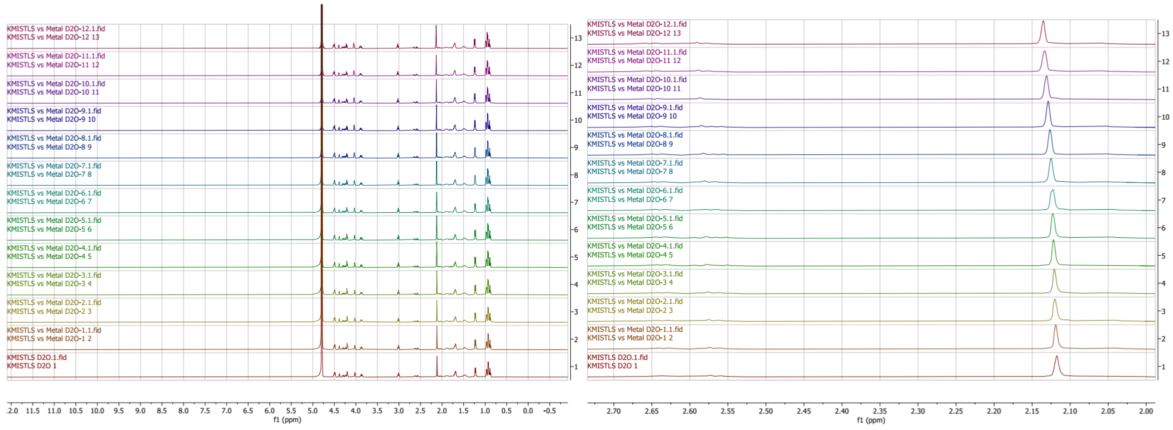


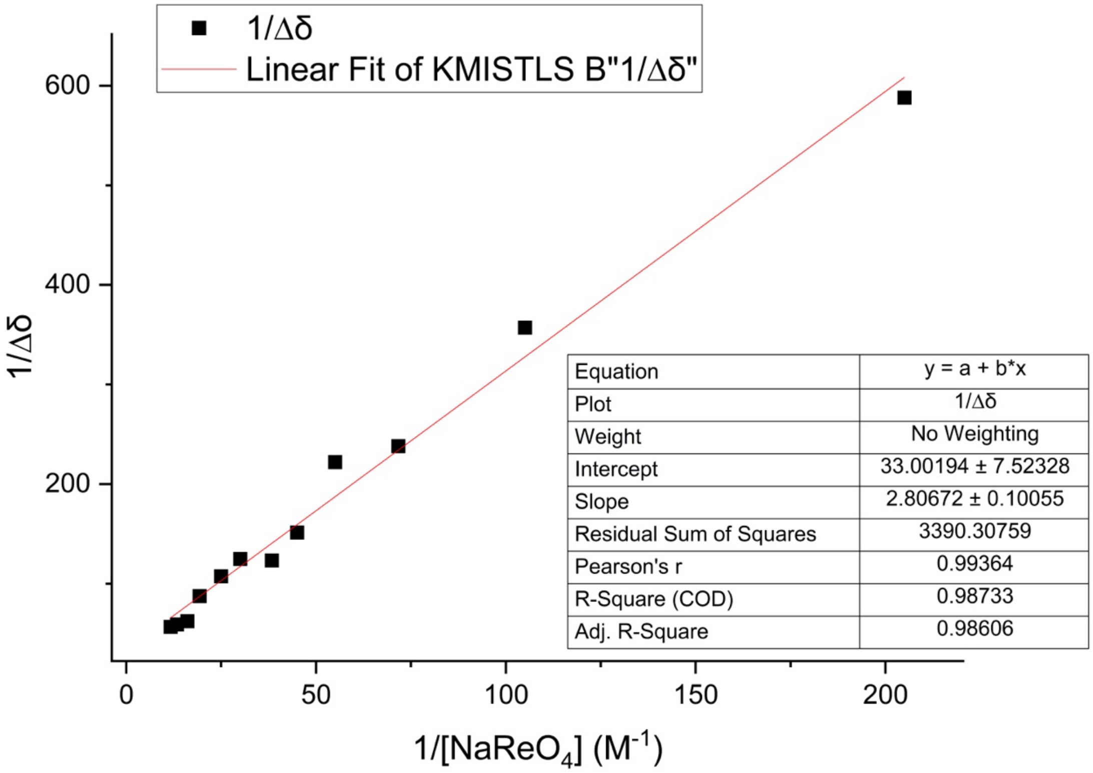

**Supplementary Figure S9.** NMR titration of ST3.

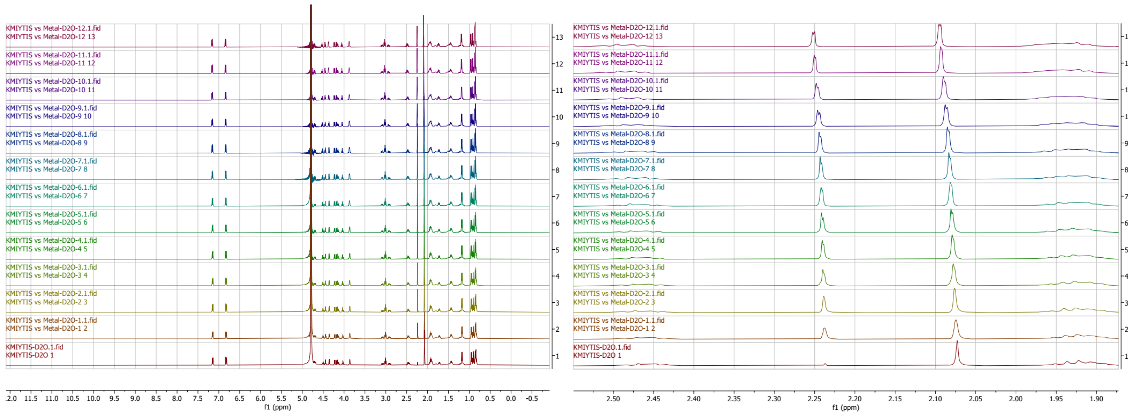

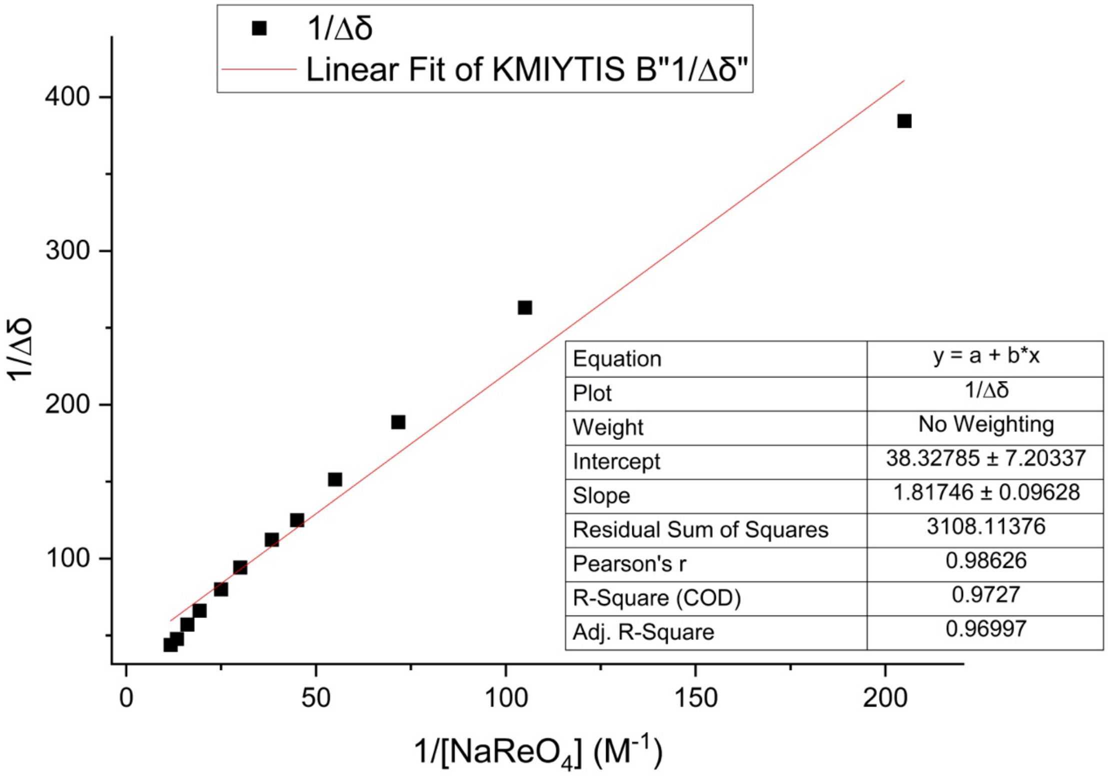

**Supplementary Figure S10.** NMR titration of ST4.

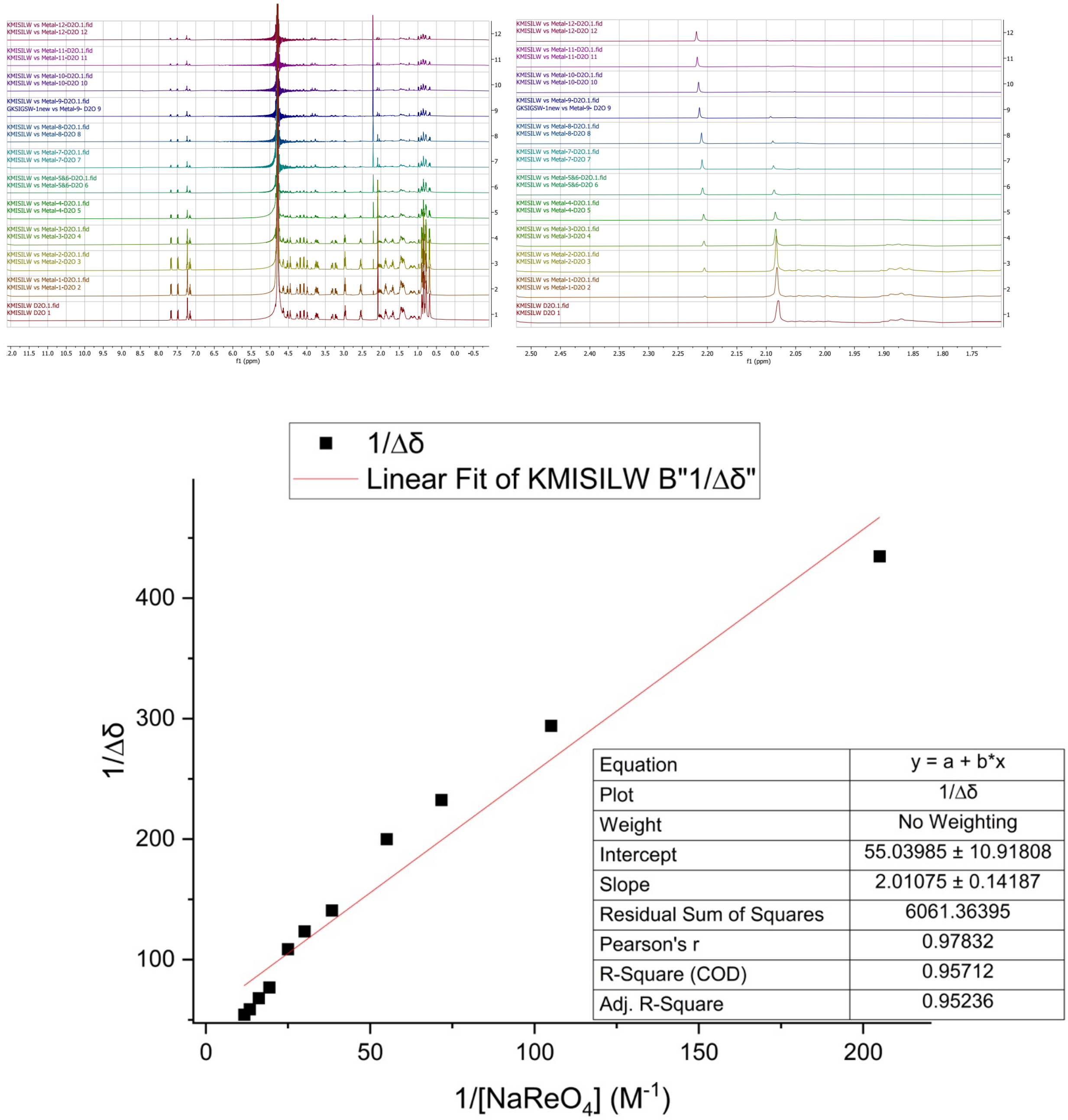


| | |
|---|---|
| Equation | y = a + b*x |
| Plot | 1/Δδ |
| Weight | No Weighting |
| Intercept | 55.03985 ± 10.91808 |
| Slope | 2.01075 ± 0.14187 |
| Residual Sum of Squares | 6061.36395 |
| Pearson's r | 0.97832 |
| R-Square (COD) | 0.95712 |
| Adj. R-Square | 0.95236 |

**Supplementary Figure S11.** NMR titration of ST5.

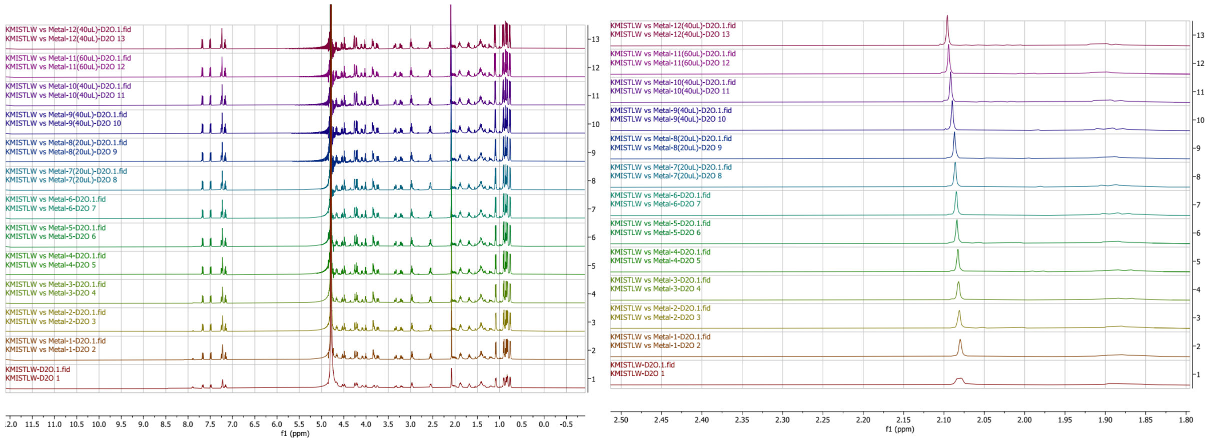


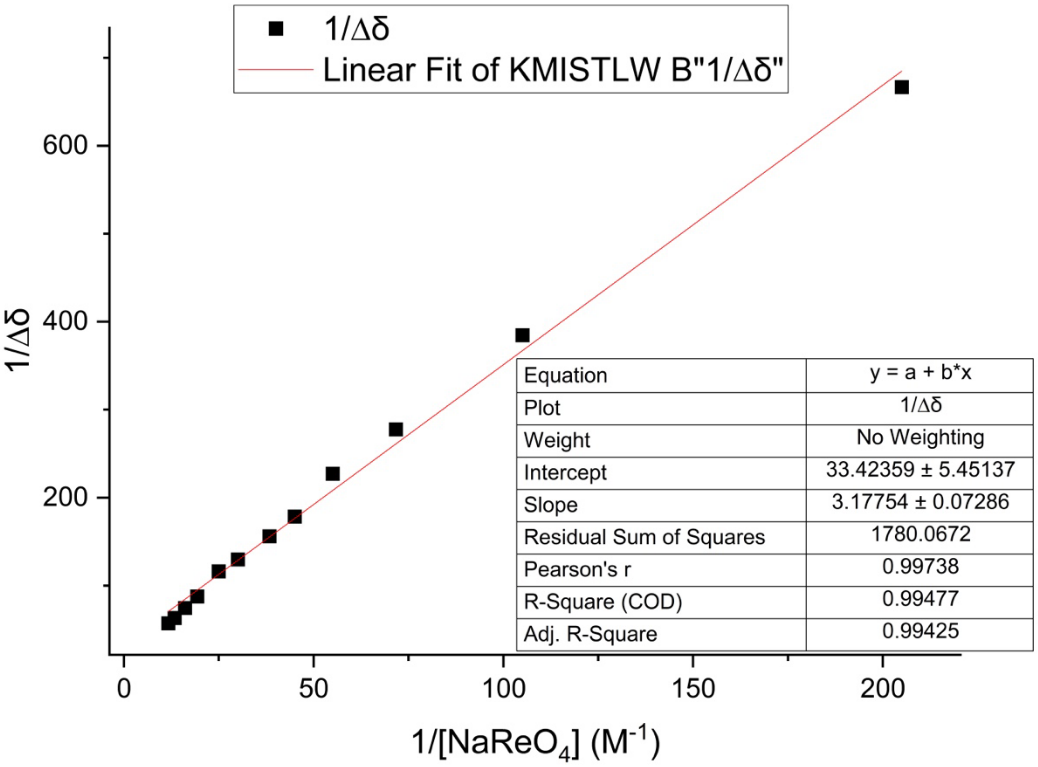

**Supplementary Figure S12.** NMR titration of ST6.

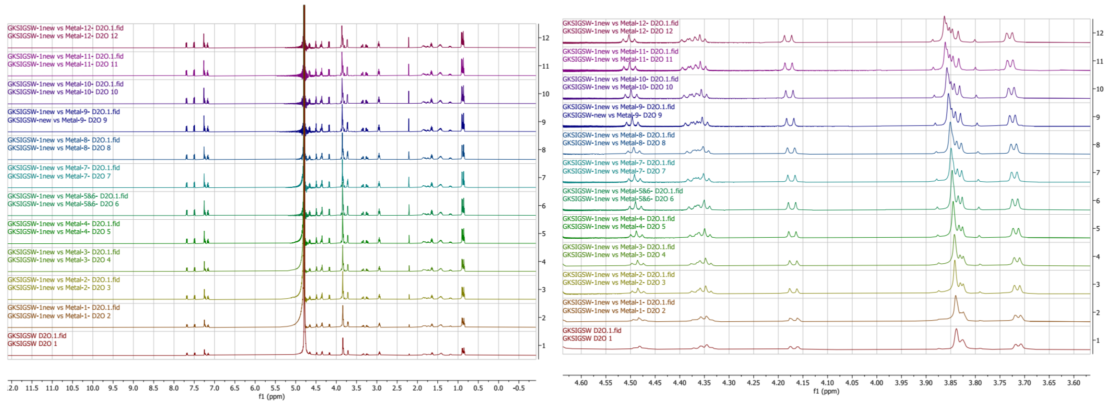


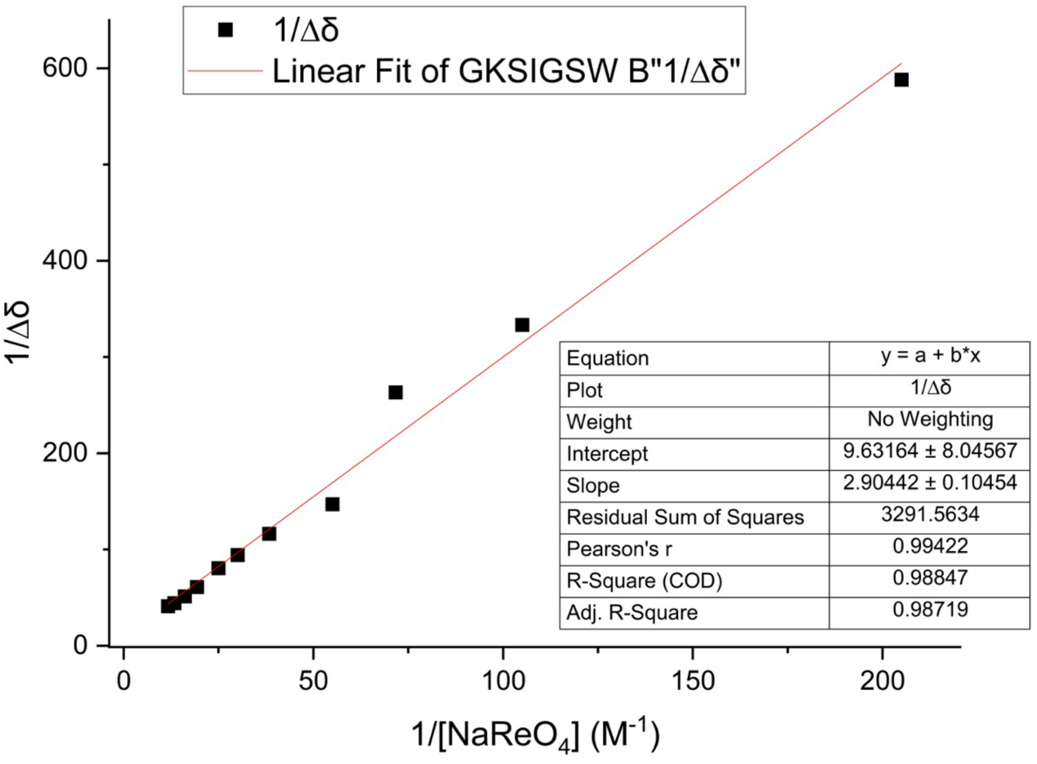


| Equation | y = a + b*x |
|---|---|
| Plot | 1/Δδ |
| Weight | No Weighting |
| Intercept | 9.63164 ± 8.04567 |
| Slope | 2.90442 ± 0.10454 |
| Residual Sum of Squares | 3291.5634 |
| Pearson's r | 0.99422 |
| R-Square (COD) | 0.98847 |
| Adj. R-Square | 0.98719 |

**Supplementary Figure S13.** Characterization of KMIS. Top) High resolution mass spectrum. [M+H] expected m/z: 477.2854, Δppm: 9. Bottom) HPLC chromatogram of purified KMIS.

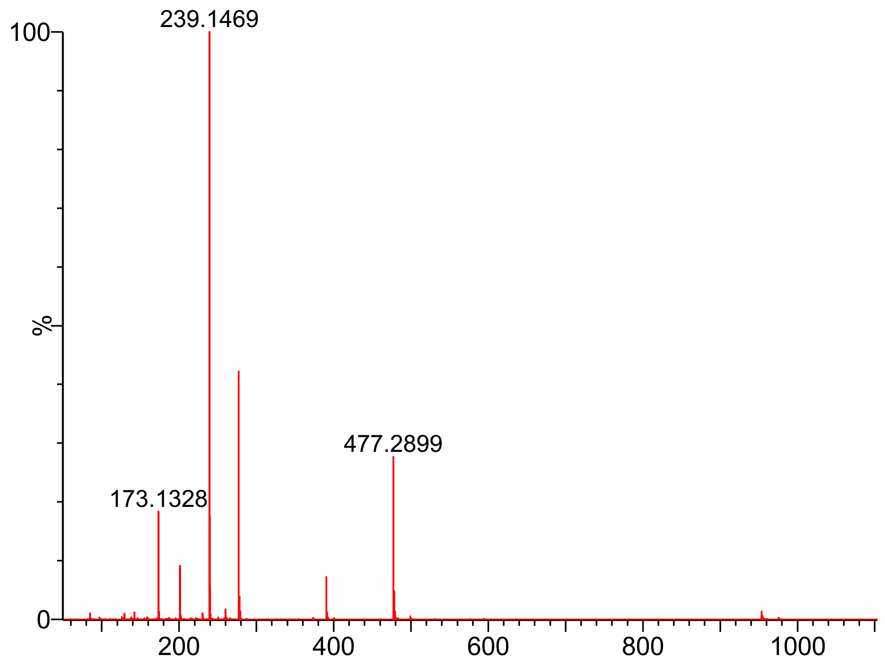


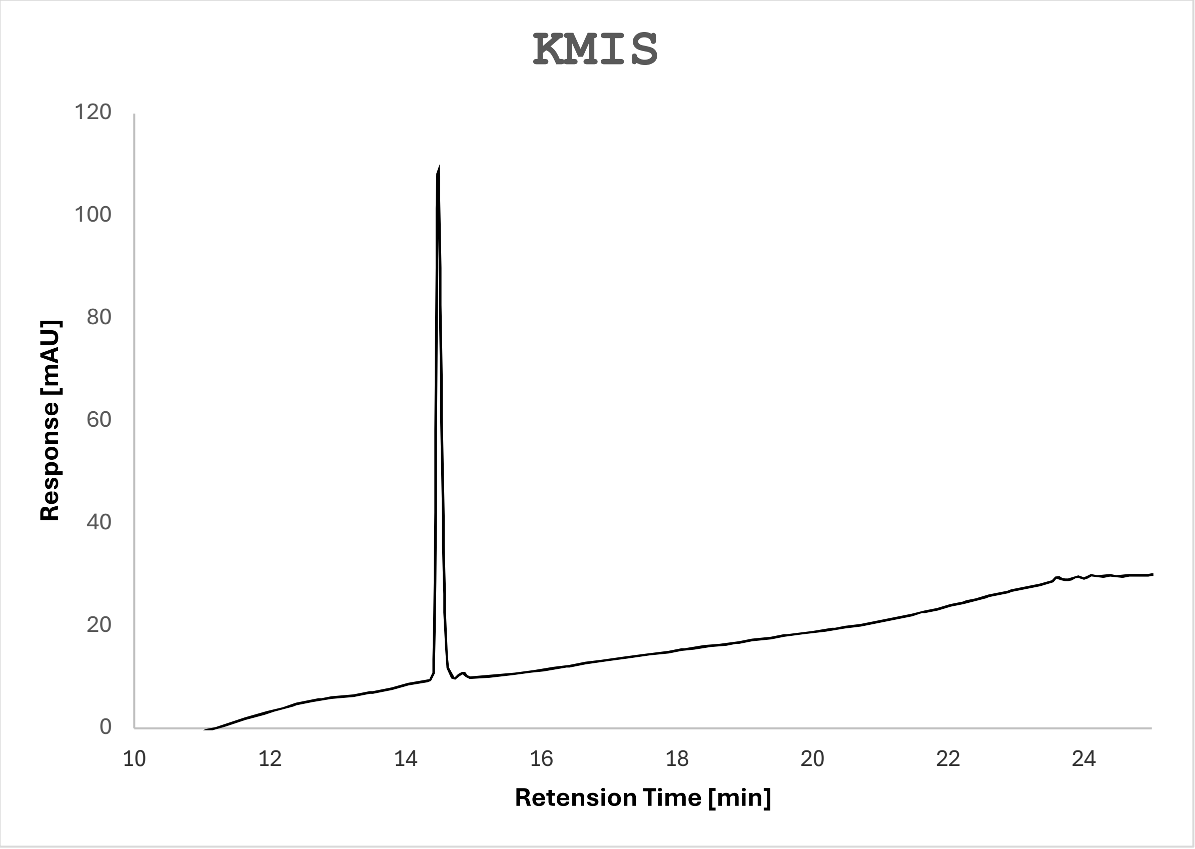

**Supplementary Figure S14.** Characterization of KMIY. Top) High resolution mass spectrum. [M+H] expected m/z: 553.3167, Δppm: 4. Bottom) HPLC chromatogram of purified KMIS.

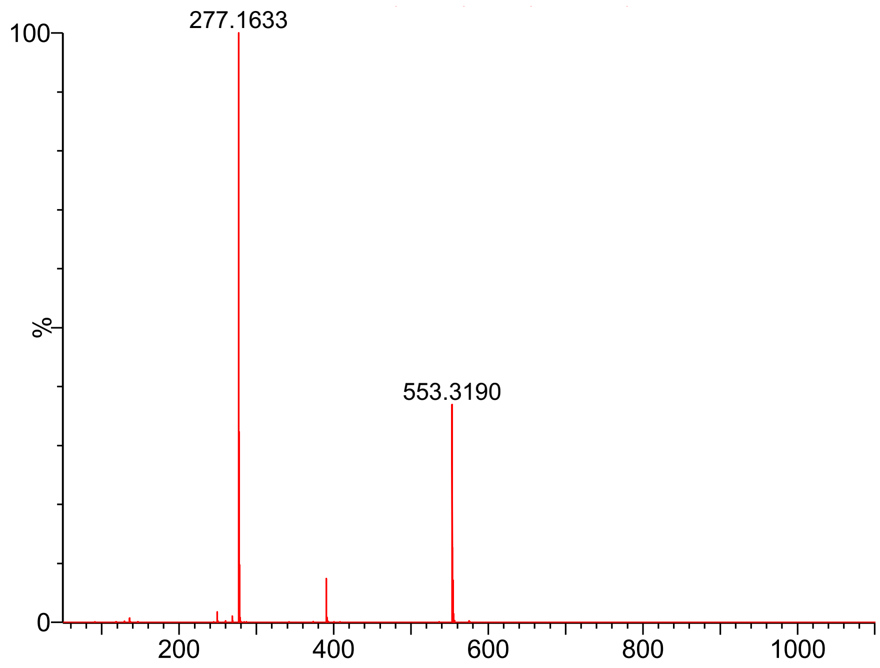


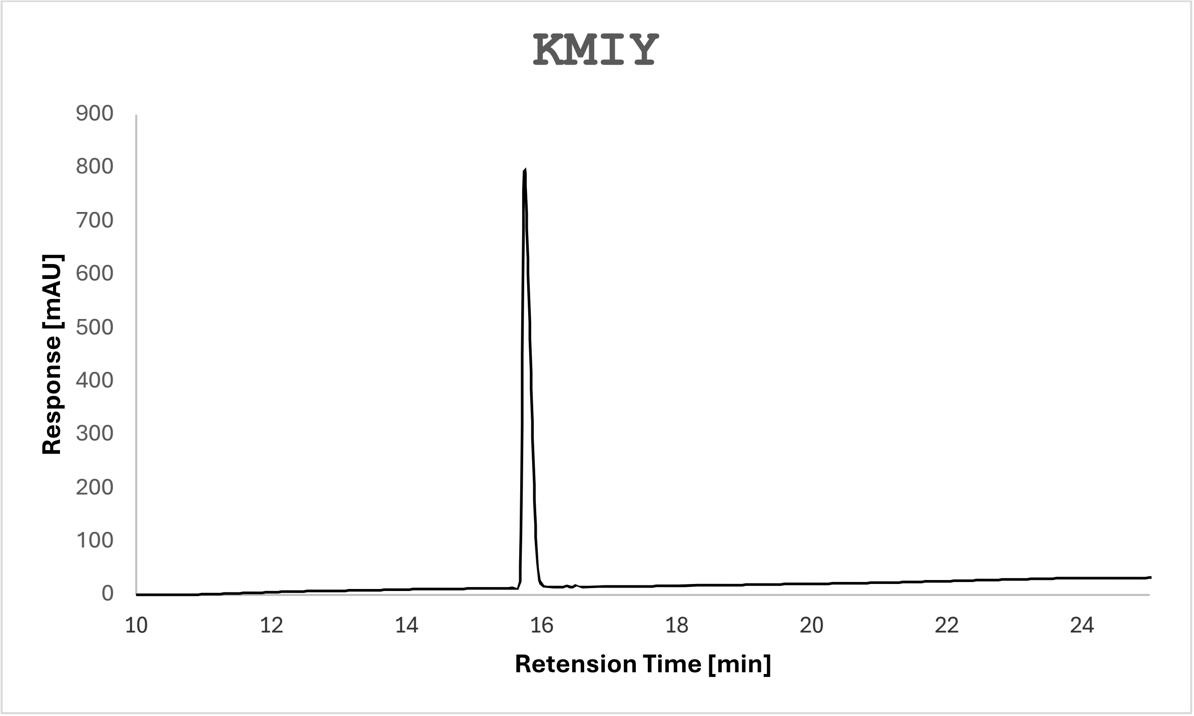

**Supplementary Figure S15.** Standard curves for metal salts at various maximum wavelengths. A) NaI at 235 nm. B) $NaNO_2$ at 235 nm. C) $NaReO_4$ at 230 nm. D) $Na_2CrO_4$ at 372 nm.

A)

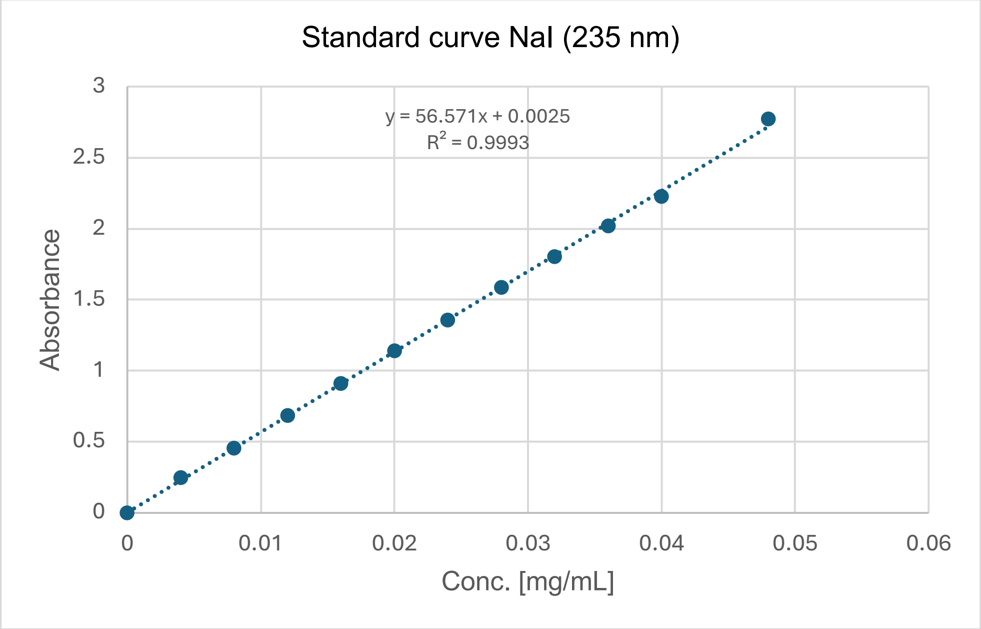


B)

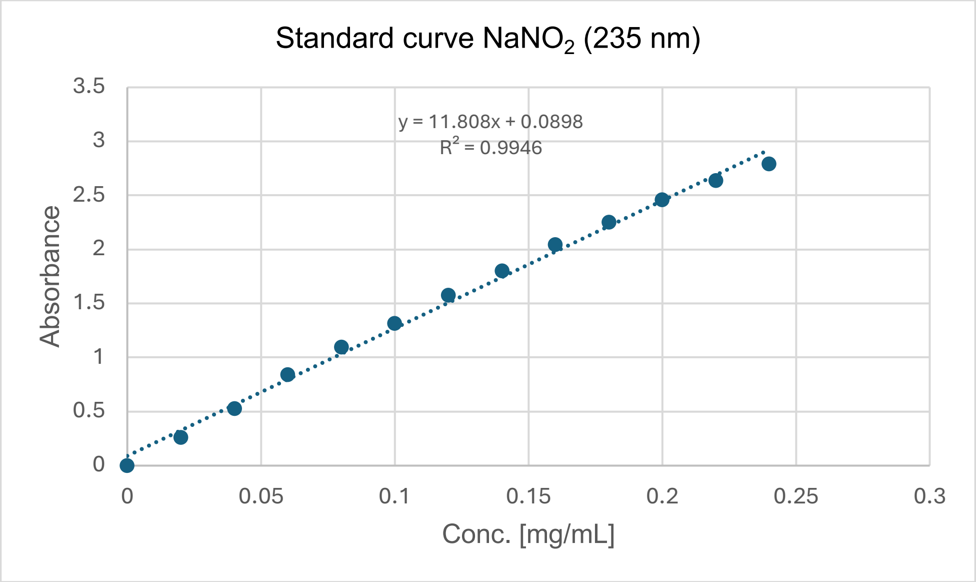

C)

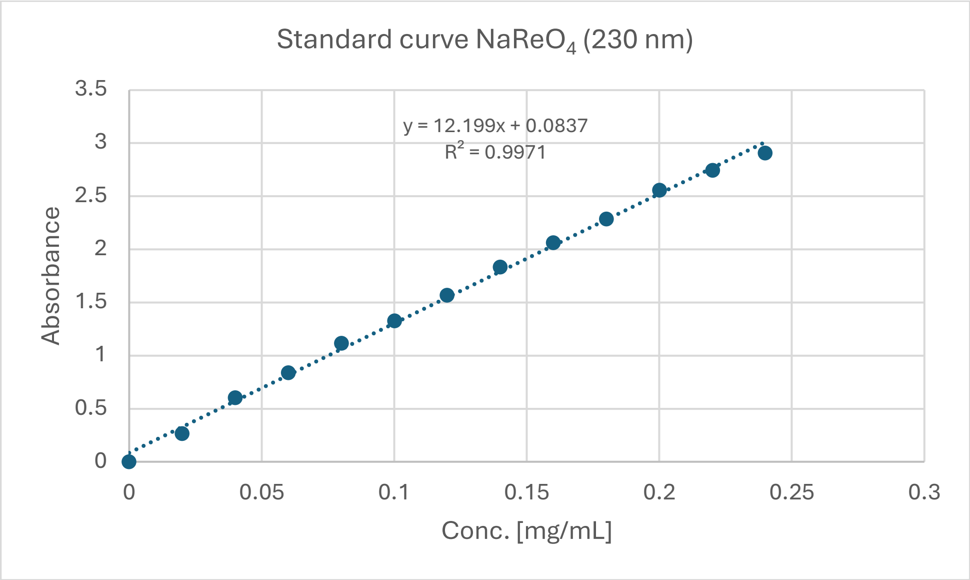
Standard curve $NaReO_4$ (230 nm)
y = 12.199x + 0.0837
$R^2$ = 0.9971
Absorbance
Conc. [mg/mL]
3.5
3
2.5
2
1.5
1
0.5
0
0
0.05
0.1
0.15
0.2
0.25
0.3

D)

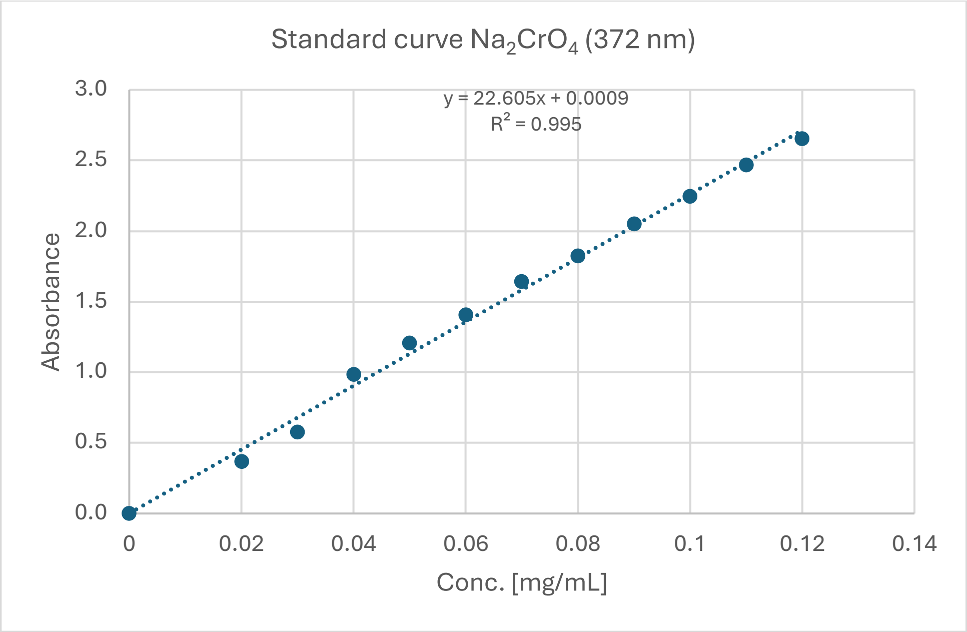
Standard curve $Na_2CrO_4$ (372 nm)
y = 22.605x + 0.0009
$R^2$ = 0.995
Absorbance
Conc. [mg/mL]
3.0
2.5
2.0
1.5
1.0
0.5
0.0
0
0.02
0.04
0.06
0.08
0.1
0.12
0.14
